# Cross-platform Bayesian optimization system for autonomous biological assay development

**DOI:** 10.1101/2021.06.23.448246

**Authors:** Sam Elder, Carleen Klumpp-Thomas, Adam Yasgar, Jameson Travers, Shayne Frebert, Kelli M. Wilson, Alexey V. Zakharov, Jayme L. Dahlin, Christoph Kreisbeck, Dennis Sheberla, Gurusingham S. Sittampalam, Alexander G. Godfrey, Anton Simeonov, Sam Michael

## Abstract

Current high-throughput screening assay optimization is often a manual and time-consuming process, even when utilizing design-of-experiment approaches. A cross-platform, Cloud-based Bayesian optimization-based algorithm was developed as part of the NCATS ASPIRE Initiative to accelerate preclinical drug discovery. A cell-free assay for papain enzymatic activity was used as proof-of-concept for biological assay development. Compared to a brute force approach that sequentially tested all 294 assay conditions to find the global optimum, the Bayesian optimization algorithm could find suitable conditions for optimal assay performance by testing only 21 assay conditions on average, with up to 20 conditions being tested simultaneously. The algorithm could achieve a seven-fold reduction in costs for lab supplies and high-throughput experimentation run-time, all while being controlled from a remote site through a secure connection. Based on this proof-of-concept, this technology is expected to be applied to more complex biological assays and automated chemistry reaction screening at NCATS, and should be transferable to other institutions.

## Introduction

There is great interest in applying artificial intelligence (AI) and machine learning (ML) to various phases of preclinical drug discovery [1]. This includes chemistry reaction screening, hit selection, and even molecular design [2, 3]. It is anticipated that AI/ML approaches can enhance the efficiency of solving many of the multifactorial problems encountered in drug discovery. This includes biological assay development, a process that can take experienced biologists months to even years to develop and validate a robust, pathophysiological relevant assay [4].

One reason for the length of assay optimization is that scientists must optimize multiple and often competing factors, with many of the variable interactions difficult to predict *a priori*. This includes the concentration of key reagents (substrate, enzymes, cofactors), temperature, buffer/media composition, timing, as well as other factors. Assay optimization can utilize several strategies including brute force, design-of-experiment (DOE), or approaches informed from historical experience or fundamental principles. Brute force approaches, which test as many variable permutations as possible, and DOE approaches, which apply statistics to systematically determine the relationship between a set of variables affecting an output, and have been used in high-throughput screening (HTS) assay optimization [5–8]. Both suffer from a lack of feedback and adaptivity. This means that they each specify the entire set of experiments to be conducted *a priori*, which means they waste resources on low-performing experiments rather than concentrating attention on high-performing experiments. Additionally, this means they each scale poorly with higher dimensions, as the number of experiments necessary increases exponentially. The cost of performing all these experiments can eventually become infeasible due to costs from reagents, time, and automated system modifications. While valuable, relying solely on historical and fundamental principles has the potential for bias, and such preconceptions about how best to optimize an assay may not be best applicable to novel biological systems.

The NCATS <u>A</u> <u>S</u>pecialized <u>P</u>latform for <u>I</u>nnovative <u>R</u>esearch <u>E</u>xploration (ASPIRE) Initiative seeks to combine advances in automated chemistry, high-throughput biological annotation, and AI/ML to accelerate preclinical drug discovery [9–11]. To assess the feasibility and utility of integrating AI/ML methods in our preclinical drug and probe discovery center, we developed an autonomous Bayesian assay optimization system. Using a facile cell-free fluorometric assay for the protease papain, we demonstrate that such an autonomous system can provide an efficient, robust set of assay conditions capable of identifying bioactive small-molecules with similar performance to conventional approaches.

## Materials and methods

### Chemicals and reagents

All general chemicals were purchased from Millipore Sigma, VWR, or ThermoFisher Scientific unless otherwise stated. Papain (from papaya latex; Sigma-Aldrich) was prepared in 50% glycerol (v/v) at a final concentration of 10 mg/mL (427 μM; 100 U/mL). The fluorogenic Z-FR-AMC dipeptide substrate (Bachem, cat # I-1160, HCl salt) was prepared as a 32.1 mM stock solution in DMSO.

### Papain cell-free enzymatic assay

Screening was performed at the NCATS screening facility [12]. The effect of test compounds on papain cell-free enzymatic activity was adapted from published protocols (**Table 1** and **Supplemental Table 1**) [13, 14]. Three multi-head liquid dispensers were configured to facilitate autonomous and continuous experiments, including a microplate washing step for microplate re-use and minimization of plastics consumption (**Supplemental Table 2**). Three critical assay design parameters, based on the NCATS *Assay Guidance Manual*, were selected for optimization: reaction time, substrate concentration, and enzyme concentration [15, 16].

**Table 1.** Standardized cell-free papain assay protocol utilized for Bayesian optimization algorithm.

| Step | Parameter | Value | Description |
| --- | --- | --- | --- |
| 1a | Reagent | 3 μL | Papain in assay buffer |
| 1b | Reagent | 3 μL | Assay buffer |
| 2 | Compound | 20 nL | Compound addition |
| 3 | Time | 15 min | RT incubation |
| 4 | Reagent | 1 μL | Substrate (Z-FR-AMC) in assay buffer addition, reaction initiation |
| 5 | Time | 15 sec, 271 g | Centrifugation |
| 6 | Detection | $(E_x/E_m) = 340/460$ nm | Fluorescence (Read 1) |
| 7 | Time | Variable | Incubation (RT) |
| 8 | Detection | $(E_x/E_m) = 340/460$ nm | Fluorescence (Read 2) |

| Step | Notes |
| --- | --- |
| 1a | Medium-binding black solid-bottom Greiner plates (789176-F). Dispense 3 μL of 1.33X papain in assay buffer (100 mM sodium acetate, pH 5.5, 5 mM DL-cysteine, 0.01% Tween-20 v/v) to columns 1, 2, 5-48 via Wako/Kalypsys dispenser. Papain was dispensed to produce 16, 8, 4, 2, 1, 0.5, 0.25, and 0.125 nM final enzyme concentrations. Optimized solution was 0.125 nM papain, final concentration. Enzyme solutions kept on ice during experiment. |
| 1b | Dispense 3 μL of assay buffer (100 mM sodium acetate, pH 5.5, 5 mM DL-cysteine, 0.01% Tween-20 v/v) to columns 3 and 4 via Wako/Kalypsys dispenser. Assay buffer solution kept on ice during experiment. |
| 2 | Compound addition (20 nL) using EDC ATS-100 acoustic dispenser. Compound solutions in source plate: 169 nM to 10 mM DMSO stock solutions. Final concentrations: 0.664 nM to 39.2 μM, respectively. Final DMSO concentration: 0.5% (v/v).<br>*Step omitted during assay optimization.* |
| 3 | Room temperature incubation. Kalypsys lids used as microplate coverings. Protected from light. |
| 4 | Dispense 1 μL of 4X substrate mixture in assay buffer (100 mM sodium acetate, pH 5.5, 5 mM DL-cysteine, 0.01% Tween-20 v/v) to columns 1-48 via Wako/Kalypsys dispenser. Z-FR-AMC substrate was dispensed to produce 16, 8, 4, 2, 1, 0.5, 0.25, and 0.125 μM final substrate concentrations. Optimized solution was 4 μM Z-FR-AMC, final concentration. Substrate solutions kept on ice during experiment. |
| 5 | Centrifugation step to remove bubbles. |
| 6 | PHERAstar optics: Excitation filter 340 nm, Emission filter 460 nm. Gain = 709. Number of flashes = 10. |
| 7 | Room temperature incubation. Incubation times 10, 15, 20, 40, 30, 45, 60, 90, and 120 min. Optimized solution was 30 min. |
| 8 | PHERAstar optics: Excitation filter 340 nm, Emission filter 460 nm. Gain = 709. Number of flashes = 10 |

### Computation

Kebotix’s Bayesian optimization software was adapted and deployed to provide suggested experimental conditions for the papain biochemical assay. Given the asynchronous quasi-batching of results, the following protocol was used: if there were any new results since the last suggestion made, and at least five results collected overall, a suggested assay protocol would be made using a sequential version of the Kebotix Bayesian optimization algorithm. Otherwise, such a suggestion would be generated randomly from the untested parameter choices. An assay optimization score was then calculated based on maximizing desirable features: sufficiently high Z’, reduction in reagent use, and shorter protocol time (**Equation 1**).

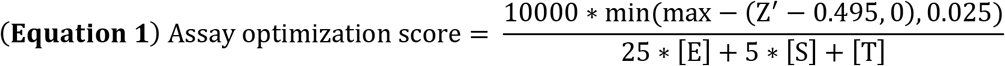

[E] represents the nanomolar concentration of enzyme; [S] represents the micromolar concentration of substrate; [T] represents the incubation time in minutes, and Z’ is a calculated metric of assay quality using the positive and neutral plate controls [17]. The denominator of the score represents a notion of the “cost” of the experiment, where smaller costs are preferred, with the coefficients on the concentrations reflecting the tradeoffs between material and time costs. The two thresholds of 0 and 0.025 applied to the quantity Z’ – 0.495 reflect the judgment that Z’ values below 0.495 are all equally undesirable, Z’ values above 0.52 (0.495 + 0.025) are all equally desirable, and those between 0.495 and 0.52 are partially desirable. Finally, the multiplicative factor of 10000 simply translated the score into the low single digits for ease of human understanding (with no effect on the optimization algorithm).

Given the results from previous assays, the Kebotix Bayesian optimization builds a model to predict the scores of untested assay conditions, with uncertainty. It then identifies the conditions that maximize a combination of the predicted score and the uncertainty in that prediction. In addition, given the functional form of the assay optimization score above, the algorithm calculates the maximum possible score for each set of conditions and focuses on conditions whose maximum possible score is higher than the current maximum, performing constrained optimization over this narrower space. This ensures that no experiments are suggested by the optimizer which are known not to improve on the current optimum.

Four variations in the experimental optimization runs were performed. Experiments within a run were started every 10 or every 20 min. For each of these intervals, a logarithmic transform was either applied or not applied before passing the options (concentrations and durations) to the optimization algorithm. The two timings were meant to illustrate performance given different tradeoffs between number of experiments and overall time spent: more frequent experiments result in fewer feedback cycles and more experiments necessary to optimize; less frequent experiments lead to more time spent on the optimization. Separately, the logarithmic transform was chosen to illustrate the value of choosing a representation of the data expected to be well-suited for optimization: The options for both concentrations and time were roughly exponentially scaled (i.e., 15, 30, 60, 120 min), and we envisioned that equally spacing these options would make for a smoother and therefore easier to optimize objective function.

### Brute force and live optimization experimental parameters

A total of 512 assay optimization scores were experimentally generated by randomly sampling each permutation (“brute force”) with the following experimental conditions: final enzyme concentration (0, 1.25, 2.5, 5, 10, 20, 40, and 80 nM), final substrate concentration (0, 0.125, 0.250, 0.500, 1, 2, 4, and 8 μM), and incubation time (5, 15, 30, 60, 120, 180, 240, and 480 min). Brute force trials with zero-value conditions (0 nM enzyme, 0 μM substrate, 0 min incubation) were used as experimental controls, but were excluded from subsequent analyses. Trials utilizing 0.250 μM substrate concentrations were excluded due to technical issues. For the Bayesian optimization live experiments, assay optimization scores were experimentally generated using the following allowable experimental conditions: final enzyme concentration (0.125, 0.25, 0.5, 1, 2, 4, 8, and 16 nM), final substrate concentration (0.125, 0.250, 0.500, 1, 2, 4, 8, and 16 μM), and incubation time (10, 15, 20, 30, 45, 60, 90, and 120 min).

### System communication

The NCATS HTS system composed of an integrated, automated robotic platform, is based upon a dynamic and asynchronous scheduling methodology in which assay plates act as the input to the system, and each plate may have associated control and compound plates as required. A method is the set of steps each assay object will complete, with each step usually associated with some peripheral device on the HTS system (e.g., liquid dispenser, compound transfer device, plate reader, etc.). To execute an assay on a microplate (“trial”), the system scheduler relies on wait queues and mutexes. At the start of an experimental run (containing multiple microplates, or trials), all assay objects are placed in a first in, first out (FIFO) wait queue. A wait queue is a queue of objects waiting for a lock, which in this case represents control of a peripheral device on the screening system. For each peripheral device, when its associated lock is unlocked, the objects acquire the lock in the order of the queue. Objects still in the queue waiting to use a locked peripheral device are in a blocked state until the lock is released. Each step, or a series of linked steps, in a method has an associated lock, with this type of resource availability-based synchronization being called a Mutex (“mutual exclusion”).

This HTS system can monitor all events occurring on the platform for each assay plate and every associated method step, can allow multiple processes to run in parallel, and can launch new processes directed from external applications. To incorporate AI/ML-driven experimentation from extramural collaborators (i.e., collaborating organizations external to the NCATS HTS facility), a messaging technique was required such that an external informatics platform could initiate experiments to be performed on the NCATS HTS system, with the resultant data then sent back for analysis to the initiating platform. To enable this messaging, a RabbitMQ server was deployed at NCATS and was made public facing to allow external messaging. Several steps were implemented to ensure data security, including the deployment of the server within the NIH firewall demilitarized zone (DMZ), the utilization of Hypertext Transfer Protocol Secure (HTTPS) communication, Transport Layer Security (TLS) protocol version 1.2, and other technical controls.

Advanced Messaging Queuing Protocol (AMQP) was used to provide a platform-agnostic method for ensuring information is safely transported between applications, among organizations, within mobile infrastructures, and across the Cloud. The screening platform can be considered as the ‘query’ portion by reading an experiment while the extramural informatics platform (Kebotix) is the ‘command’ portion issuing new experiments to be performed. In our system, this informatics platform produces a message that routes through the exchange to be placed within the experimental queue at the HTS site. This message initiates an experiment, and consists of a unique identifier for experiment tracking, and in this specific report, details on the desired enzyme concentration, substrate concentration and incubation time. Using LabView [18], the HTS system consumes this message from an experimental queue and generates a method specific to it based upon the experimental conditions requested to be programmatically launched as an assay on the platform. During the experiment, any plate read steps that generate data trigger the HTS system to generate a new message (containing the unique identifier and resultant data) that is added to the results queue. Upon completion of an experiment, another message is produced by the HTS system and sent to the results queue to let the informatics platform know that the experiment is done. These messages in the results queue are then consumed by the informatics platform such that data generated by the experiments run on the HTS system can be analyzed and any respective models used to generate new experiments can be updated. The informatics platform could then programmatically initiate new experiments by producing another message to be sent to the experiment queue. This process would complete until some stop criterion is met. The entire system is event-driven, with commands issued by Kebotix to initiate a new experiment upon demand (assuming there are resources available to perform the request). Notably, an asynchronous operation feature was incorporated in which synchronization is not based upon time, but rather upon resource availability. This allows the scheduler to run multiple experiments in parallel in an asynchronous fashion. This means that multiple commands can be issued and multiple queries can be in process simultaneously without being time dependent as the duration of a particular experiment or time to process the results and make new predictions will be variable.

### Data simulations

Simulations were performed using the brute force data collected to identify typical behavior of variations on the optimization algorithm. In these simulations, the timings associated with the experimental process were mimicked, with the results of each suggestion being drawn from the brute force data upon completion. The following ten variations were considered: new experiments started every 10, 20, 30, 60 min or sequentially, and with and without the logarithmic transform applied to the options. Two scenarios were considered. In the first (‘trial simulations’), the optimization would proceed until the optimum conditions were suggested, immediately stopping, and approaches would be assessed by how many experiments they required to find the optimum. In the second (‘budget simulations’), the optimization would proceed for the designated budget of 40 trials and then stopped, and variations would be assessed according to whether the optimum was discovered and whether it was discovered within the first half of the budget (20 trials). In each of these scenarios, and for each of the 10 variations considered, 1,000 simulations were run.

### qHTS data analysis and statistics

Reference papain inhibitors (**Supplemental Table 3**) were tested in 11-point serial three-fold titration (0.664 nM to 39.2 μM final concentrations). The Library Of Pharmacologically Active Compounds (LOPAC^1280^) was tested in qHTS format in 7-point serial four-fold titration (31 nM to 50 μM final concentrations) [19]. Data from each assay were normalized to intra-plate controls (neutral control, DMSO; positive control as noted). The fluorescence intensity difference between Read 2 (post-reaction) and Read 1 (shortly after substrate addition) was used to compute reaction progress. The same controls were used for the calculation of the Z’ factor. Concentration-response curves (CRCs) were fitted and classified as described previously. IC_50_ values were calculated using Prism software (version 9.1.0, GraphPad), sigmoidal dose-response (variable slope). Data was curated using the Palantir Technologies (Washington, DC) data integration Foundry platform (NIH Integrated Data Analysis Platform, NIDAP), which is configured to ingest all HTS results generated at NCATS and harmonized this data with other sources such as ChEMBL and OrthoMCL. All qHTS screening results are publicly available at PubChem (AIDs 1645873, 1645872). The chemical structures were standardized using the LyChI (Layered Chemical Identifier) program (version 20141028, github.com/ncats/lychi). *P*-values were one-sided and used a Bonferroni correction where applicable.

## Results

### Assay and platform overview

A cell-free assay for papain protease activity was chosen for proof-of-concept. This assay was ideal because its low cost and simplicity is suitable for many experiments (“brute force”), and previous NCATS experience would allow for historical comparisons [13, 14]. In this fluorescence intensity assay, papain enzymatically cleaves the quenched AMC fluorophore from a dipeptide substrate to generate non-quenched free AMC fluorophore whose fluorescence intensity is proportional to papain activity in the absence of compound-mediated interferences (**Figure 1**).

**Figure 1.**
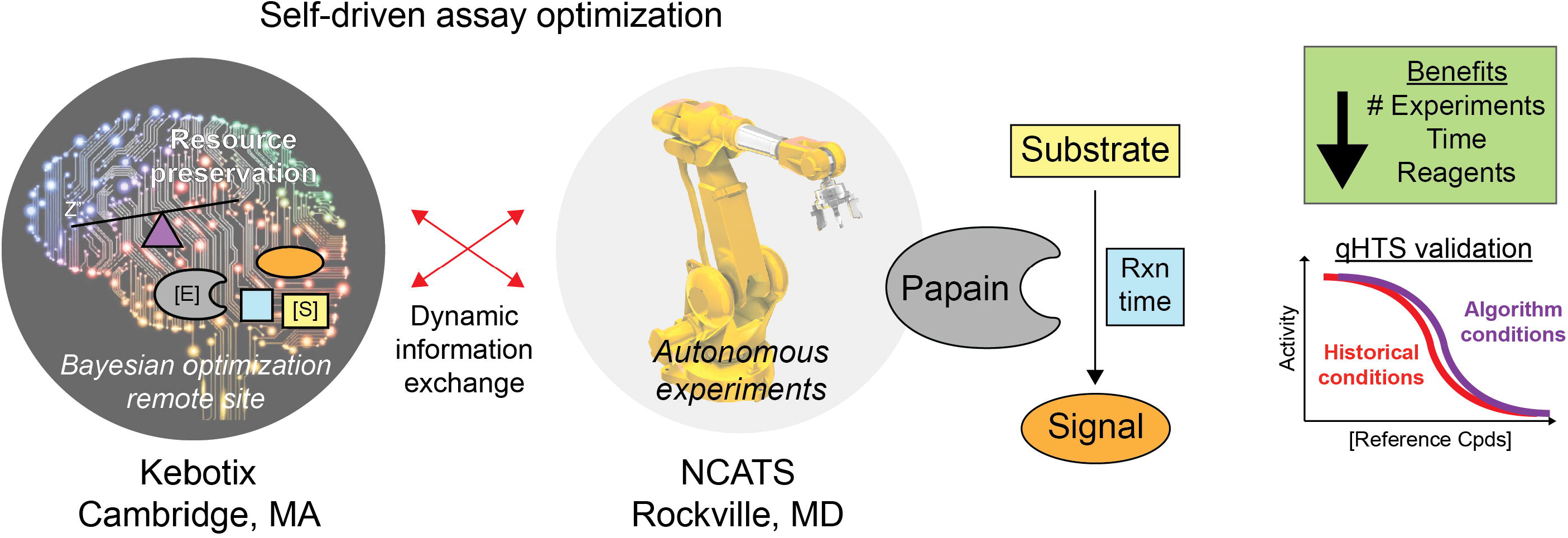
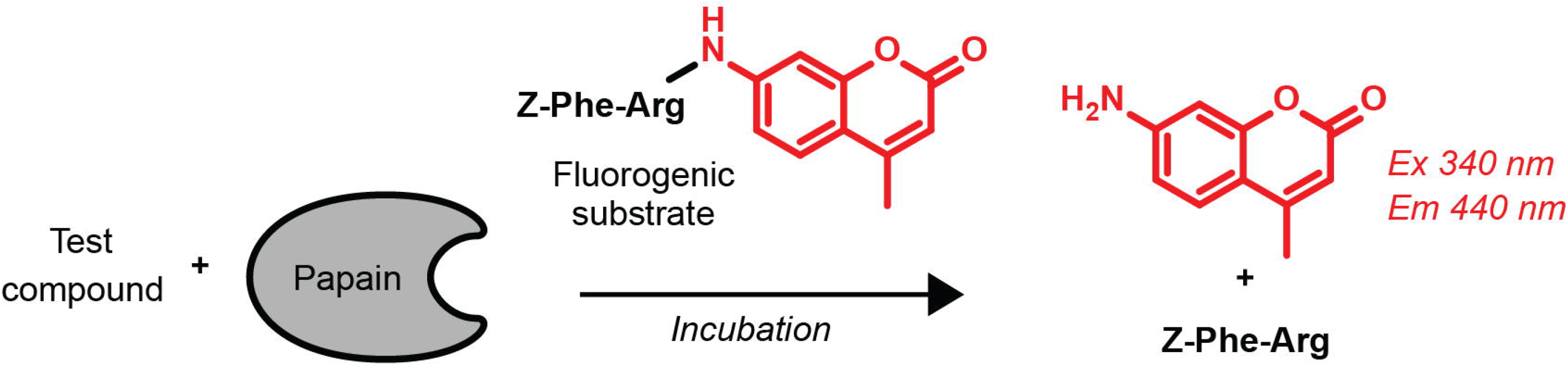
Papain assay schematic. Papain hydrolyses the AMC fluorophore from the dipeptide substrate, which leads to an increase in blue fluorescence. Inhibitors of papain activity are expected to decrease the fluorescence intensity readout. See also **Table 1** and **Supplemental Table 1** for standardized protocols of algorithm-derived and historical protocols, respectively.

An AMQP method was developed to support secure, platform-agnostic information sharing between applications, among organizations, within mobile infrastructures, and across the Cloud (**Figure 2**). The entire system is event-driven, with commands issued by a command site (i.e., Kebotix) initiating a new experiment on-demand and asynchronously. This latter feature allows controllers to run multiple trials in parallel if resources allow.

**Figure 2.**
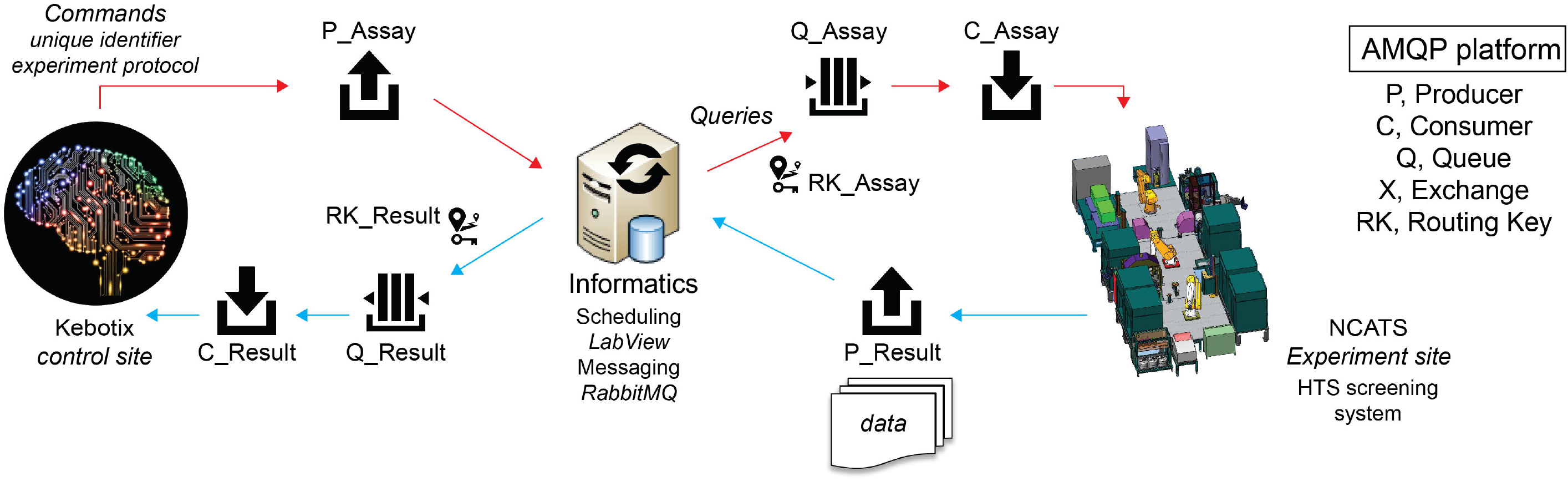
Design of autonomous assay optimization platform. The system utilizes Advanced Messaging Queuing Protocol (AMQP) to facilitate communication between the NCATS HTS system, a shared informatics resource, and the Kebotix extramural Bayesian optimization command platform. Producer (P): an application that sends messages; Queue (Q): a buffer that stores messages; Consumer (C): an application that receives messages; Exchange (X): a router that receives messages from P and pushes them to C; Routing Key (RK): a relationship that specifies that a Q is interested in messages from an X.

### Brute force biological assay optimization

To help assess the effectiveness of the Bayesian assay optimization approach, the assay optimization parameters were tested sequentially using a brute force approach. Notably, this was a completely autonomous process that required no human intervention, and allowed for multiple other independent projects to be run on the same testing platform throughout the duration of the experiment. Testing each permutation required 512 individual experiments and approximately 96 h of automated experimentation time (**Figure 3A**). Based on the assay optimization scores (**Equation 1**), the optimal experimental conditions were: variable substrate concentrations (low μM), low nanomolar enzyme concentrations (1.25 to 2.5 nM), and intermediate incubation times (15 to 30 min; **Figure 3B**).

**Figure 3.**
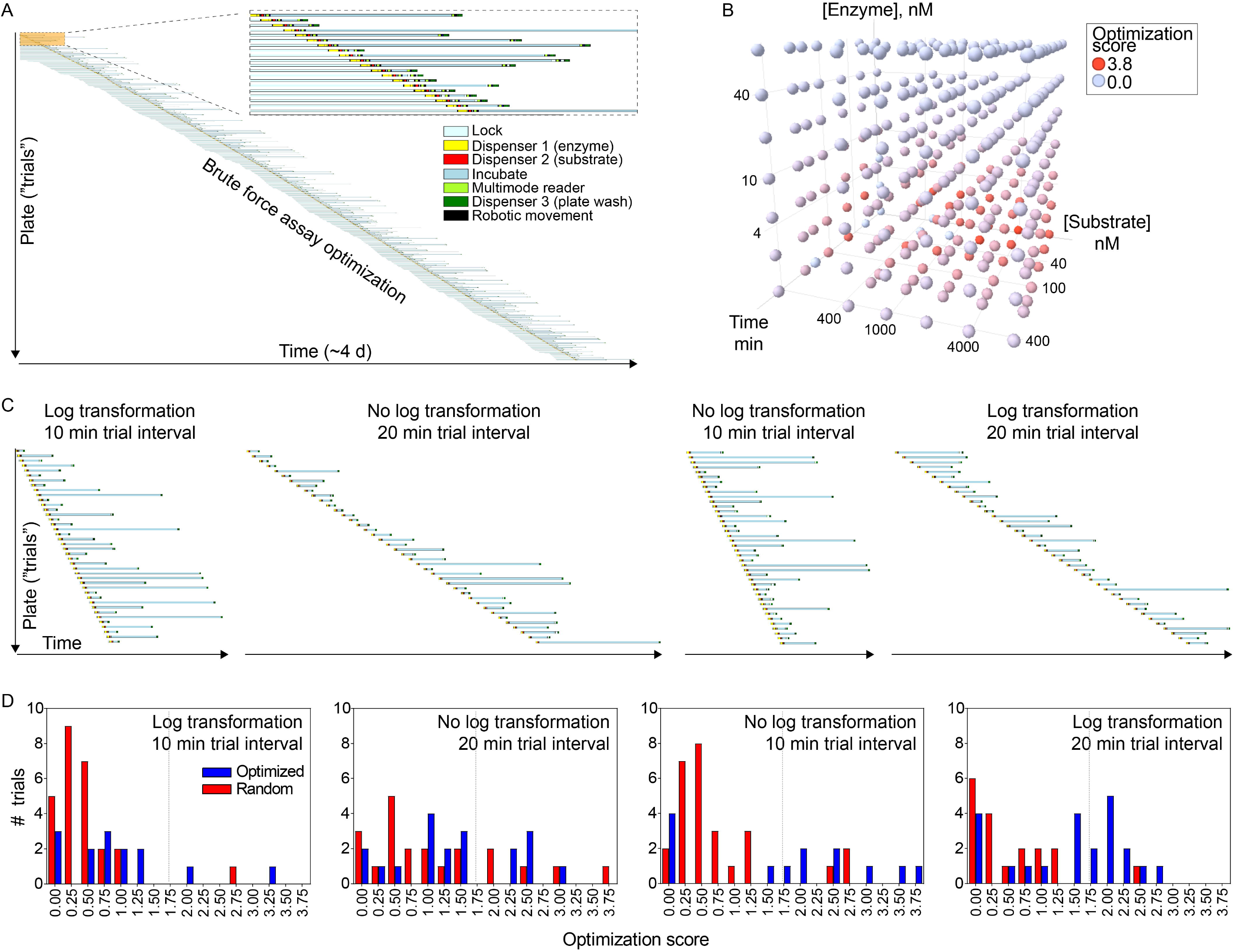
Optimization of papain biochemical assay by brute force and Bayesian optimization approaches. **(A)** A brute force approach determined optimal papain assay conditions by randomly testing each permutation of substrate concentration, enzyme concentration, and incubation time. Gantt chart of the brute force approach demonstrates the autonomous, asynchronous, event-driven system can function in a continuous fashion for over four days. (**B**) Summary of assay optimization for the brute force, scored by weighting time, Z’, and reagent consumption. (**C**) Gantt charts of the Bayesian optimization approach, performed in four independent runs testing each combination of slow/fast experiment queuing, and log-transformed/non-log-transformed data. (**D**) Distribution of experiment quality from panel (**C**) for non-transformed and log-transformed data, and slow and fast experiment queues.

### Bayesian assay optimization

The Kebotix Bayesian optimization algorithm was used to determine the optimal assay conditions while blinded to the brute force results. To facilitate future improvements in the algorithm for assay optimization, the algorithm was performed in four independent runs each with a unique combination of parameter transformations (log or non-log) and interval time between trials (10 or 20 min). Not surprisingly, the runs with increased trial intervals required more time compared to the runs with shorter trial intervals (**Figure 3C**).

Live optimization runs illustrate the performance of the optimization algorithm in action. While the true optimum is unknown without all suggestions having been attempted, a simple comparison of the performance of the randomly generated suggestions and those generated by the optimum can illustrate the performance of the optimizer (**Figure 3D**). Across the four runs, optimized suggestions achieved assay optimization scores above the arbitrary threshold of 1.4 in 34 of 68 (50%) attempts, while random suggestions achieved optimization scores above 1.4 in only 12 of 92 (13%) attempts, four-fold less frequently. Even accounting for the arbitrariness of this threshold, this difference is significant (*p* = 3 × 10^−6^). This demonstrates the optimizer can enrich for high quality experiments and provide biologists with more viable choices for selecting final conditions. This might be advantageous in cases where expert knowledge is to be incorporated that is difficult to incorporate into an objective function.

### Further validation by simulation

Ground-truth data generated by the brute force approach enabled further validation by simulation of the overall platform, as well as individual components of the adaptive learning processes itself like logarithmic data transformation and the timing of experimental queues. Two types of simulations were performed: trial simulations and budget simulations. Trial simulations assessed the number of assay condition trials needed to identify the optimal assay condition (allowing up to the full 294 trials), while budget simulations assessed how often a run could identify the optimal assay condition with 40 or less trials.

In trial simulations, variations of the optimizer both with and without the logarithmic transform found the optimum in only 11 to 21 trials on average, a 7- to 13-fold improvement on brute force, with the expected variation according to the frequency of experiments (**Figure 4A**). This illustrates the tradeoff between time and experiment costs: Running more frequent experiments (shorter intervals between consecutive trials) saves time, while running less frequent experiments requires fewer to find the optimum. The improvement using log transform was minimal (mean difference less than 1 experiment), indicating the transform was not important for the algorithm to find the optimum.

**Figure 4.**
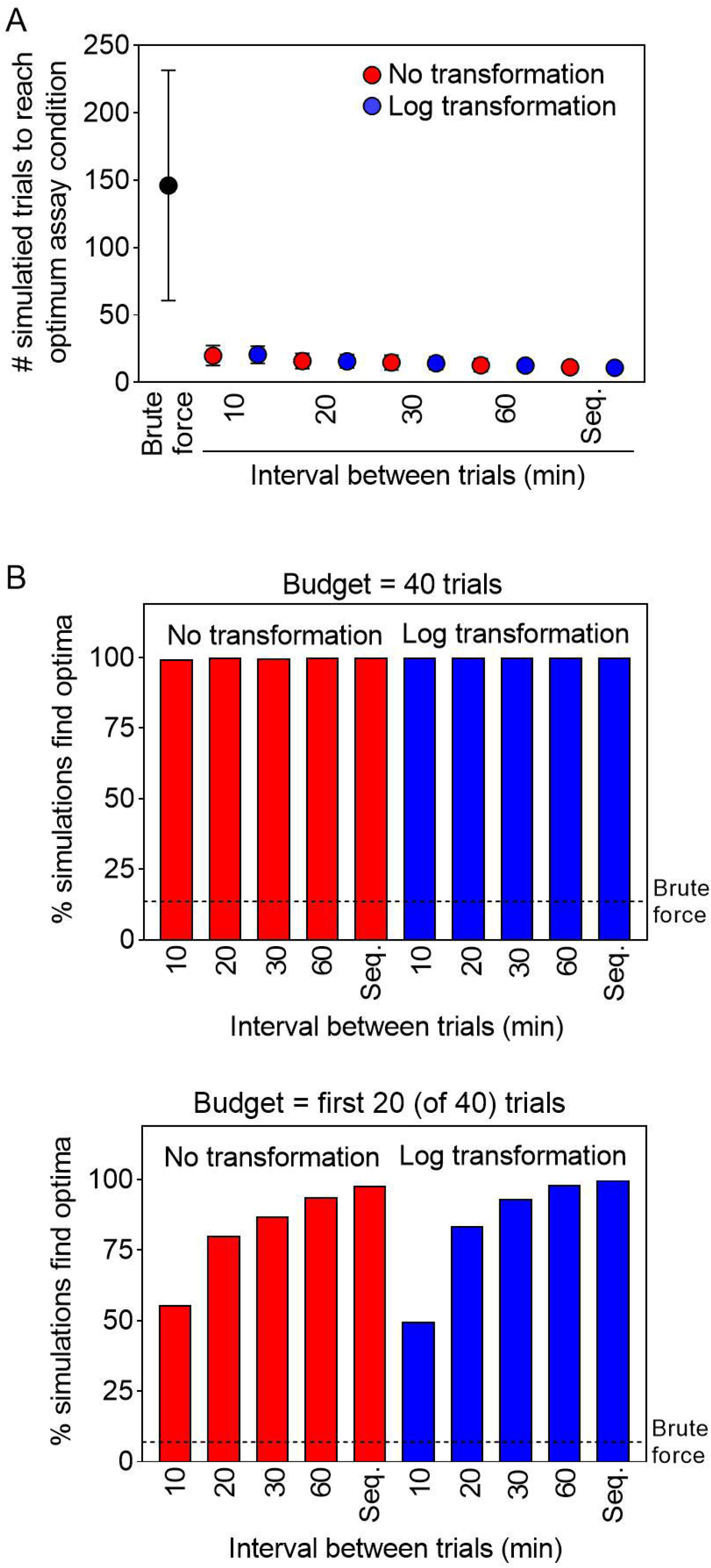
Validation of adaptive learning and optimization processes using brute force-derived simulation data. Simulations compared adaptive learning to brute force, and the effects of time delays between subsequent experiments for non-transformed and log-transformed data. (**A**) Trial simulations for runs until the assay condition optimum was found (up to 294 trials per simulation run). Data are arithmetic mean ± SD. (**B**) Top; budget-based simulations (n = 1000) where each simulated run was allotted up to 40 trials. Bottom; analysis of first 20 trials.

In budget-based simulations where each run was allotted up to 40 trials, the optimization algorithm was able to identify the optimum at least 99.4% of the time in all variations, seven-fold more frequently than the brute force approach with *p*-values less than 10^−846^ (**Figure 4B**). When simulating a more stringent budget (20 trials), all variations were able to identify the optimum at least 49.7 of the time, and as frequently as 99.8% for sequential trials applying the logarithmic transform, offering a similar 7- to 15-fold improvement on brute force (**Figure 4B**). In both simulation formats, extending the time between experiments beyond ten minutes decreased the number of experiments needed to find the optimum (**Figure 4**). These simulations demonstrate the efficiencies of adaptive learning for assay optimization, and should help in future experiments with this platform.

### Validation of optimized conditions with reference small molecules and pilot qHTS

Select compounds were tested for inhibition of papain enzymatic activity to further assess the robustness and applicability of the algorithm-derived solutions. At this stage, an additional compound transfer step was added to the assay protocol, and CRCs were automatically analyzed by an in-house informatics platform. A series of nearly two dozen reference papain inhibitors spanning nanomolar to micromolar potencies was selected based on in-house data from a papain counter-screen used during a qHTS campaign targeting the protease cruzain [13, 14]. These reference compounds were tested using the historical counter-screen conditions (**Supplemental Table 1**) and the optimal conditions determined from the algorithm. While there was some systematic bias observed (Bland-Altman bias = 0.3 ± 0.2), there was gross concordance for the IC50 values determined using the historical conditions and those identified by the algorithm (r^2^ = 0.93; **Figure 5A**). Notably, there were some differences in CRC shape between the two assay conditions (**Figure 5B**).

**Figure 5.**
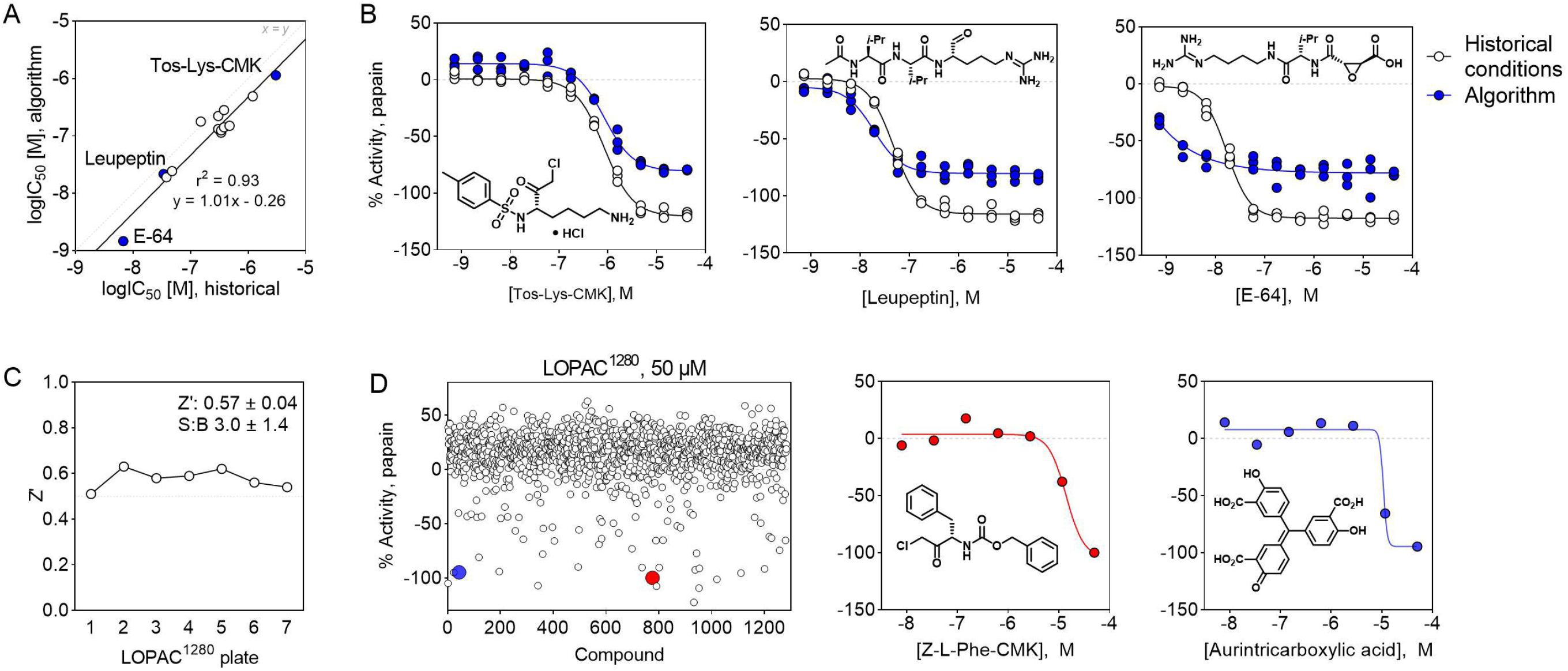
Validation of the algorithm-optimized experimental conditions for a cell-free papain inhibition assay by small molecules. (**A**) Papain inhibition by a reference set of small-molecule papain inhibitors shows general agreement between historical assay conditions and the algorithm-derived conditions. (**B**) Representative concentration-response curves of reference papain inhibitors. Data are three technical replicates from one of three independent experiments. (**C**) The algorithm-derived conditions showed acceptable assay quality metrics using the LOPAC^1280^ in qHTS format. Shown are representative data from one of six independent experiments. (**D**) The algorithm could identify small-molecule inhibitors of papain enzymatic activity using the LOPAC^1280^ in qHTS format. Right; representative concentration-response curves of papain inhibitors identified in the LOPAC^1280^ qHTS. Shown are representative data from one of six independent experiments.

The applicability of algorithm-optimized assay conditions was also assessed by a pilot qHTS using the LOPAC^1280^. The Kebotix conditions showed acceptable assay quality metrics using the LOPAC^1280^ in qHTS format (**Figure 5C**). This autonomously optimized method could identify putative small-molecule inhibitors of papain enzymatic activity using the LOPAC^1280^ in qHTS format, including a likely covalent inhibitor Z-L-Phe-chloromethyl ketone and the prototypical nuisance compound aurintricarboxylic acid (**Figure 5D**). As the pilot qHTS was intended as proof-of-concept, additional characterization of primary hits was not pursued. Such experiments would ordinarily include orthogonal assays and counter-screens for common compound interferences such as light interference, nonspecific electrophiles, and aggregators, as outlined in the NCATS *Assay Guidance Manual* [15, 16].

## Discussion

We developed a cross-platform, Bayesian optimization system and applied it to autonomous biological assay optimization. As proof-of-concept, the system was able to efficiently optimize a papain biochemical enzymatic assay. The optimized assay conditions were then validated by testing a collection of reference papain inhibitors and a pilot qHTS library. These assay conditions provided practical solutions, as testing with the AI/ML-derived assay conditions recapitulated the performance of inhibition data from historical protocols, and could identify a series of inhibitions amongst the LOPAC^1280^.

Several notable features can be achieved by utilization of the developed platform. First, the proposed approach can function continuously without human supervision once reagents were prepared and system checks were performed. Second, by applying the asynchronous, event-driven method, the screening facility was able to perform other tasks on unrelated projects at the same time, including other qHTS campaigns. Third, the unique messaging system implemented in the proposed platform allows to run the physical experiments in one place (Rockville, MD, USA) while being controlled by an off-site location (Cambridge, MA, USA) through a secure electronic connection. This last feature may be especially beneficial in situations with limited on-site availability (e.g., pandemic-related restrictions).

There are several advantages of this Bayesian optimization methodology relative to traditional and brute force approaches. These algorithmic approaches can be more efficient than traditional assay optimization approaches such as brute force or DOE, resulting in increases in operational efficiency and cost savings. This is especially important for expensive reagents where the number of optimization experiments is cost prohibitive, and precious reagents such as difficult to expand or rare cell lines. Even for cheap experiments, they also do not require the involvement of the researcher in the optimization run itself, which provides additional savings when paired, as in this example, with an autonomous experimentation platform. Finally, while not demonstrated in this work, they offer a better scaling than traditional methods at high dimensions, which also present additional challenges for humans to analyze.

The Bayesian optimization approach also had some notable limitations, though many of these limitations can likely be overcome in future iterations. We can distinguish two types of limitations: Those which were necessary for achieving the full comparison to demonstrate performance of the Bayesian optimization approach relative to brute force, and those which would apply even without the need to run the brute force experiments as a comparison.

For examples of the first type, it was necessary to only optimize across a small number of dimensions (3), to allow the brute force experiments to all be run in a reasonable time frame. That exhaustive search was only necessary in this case to provide an instance of the objective function on the complete optimization space to allow for simulations to definitively demonstrate the improved performance of the algorithm. In real use, we would expect a larger number of parameters to be varied without the need to exhaustively attempt all possible combinations merely for simulation purposes. A second example of this type is the limitation of the options to particular choices. For instance, nothing about the experiment necessitates that “30 min” and “60 min” are possible durations but “45 min” is not. We chose to limit the options to these choices to allow the brute force to exhaustively test all possibilities, but in actual use, duration could be a continuous variable with only lower and upper limits defined.

One limitation of the second type is that the varying concentrations of substrate and enzyme were achieved by pre-mixing the batches of each at the selected concentrations. This aided in the development, but during the optimization run, some concentrations will tend to be chosen more frequently than others, resulting in wastage. This limitation could be overcome with dynamic dilution approaches, which would also allow concentrations to be varied continuously, as with duration as discussed above. A final limitation of the second type is that the space of all possibilities must be defined at the outset of the optimization. This is shared with other traditional approaches like DOE, but more manual optimization approaches may choose to vary a different set of parameters midway through the run based on the results thus far.

Future work will focus on applying this technology to more complex biological systems such as multistep biochemical assays and cellular assays. Additional optimization parameters can include variables such as reaction solution composition (buffer, pH, detergent, chelation agents, reducing agents, salts, cofactors, decoy proteins), key reagents (enzyme batches and sources, antibodies, cell lines), reaction conditions (temperature), and processing steps (washes, reader settings), amongst others. The assay optimization score can also be improved or tailored to specific applications. For example, while extremely useful for many assays, the Z’ metric threshold of 0.5 may not be suitable for certain assays [20].

We hypothesize this approach could be rendered more efficient for assay optimization by including multiple experimental conditions per assay plate. If the optimization score can be satisfactorily estimated using only a portion of the wells on a given plate, multiple conditions could be combined on the same plate, although some factors (e.g., time) would need to be shared. Adapting this platform to reaction screening is another future endeavor, especially in the context of NCATS ASPIRE Initiative which includes a significant automated chemistry component. Other potential applications of this technology could involve hit selection for confirmatory testing and in iterative screening approaches. It is therefore anticipated that the continued development of this integrated, AI-driven autonomous system will enhance the efficiency of assay development and other complex tasks in early preclinical drug discovery at NCATS. Through dissemination of lessons learnt and best-practices, the scientific community at-large should also benefit.

## Author contributions (CRediT)

Conceptualization: SE, SM, CK, DS. Data curation: KMW. Formal analysis: SE, AY, JLD, CKT, SM. Investigation: SE, AY, CKT, JT, SF. Methodology: SE, SM. Software: SE, DS, SM, AVZ. Supervision: AGG, GSS, AS, SM. Validation: AY, CKT. Visualization: JLD, SE, SM. Writing, original draft: JLD. Writing, review and editing: all authors.

## Acknowledgements

The authors acknowledge the following financial support: intramural program of NCATS and NIH. The content of this publication does not necessarily reflect the views or policies of the Department of Health and Human Services, nor does mention of trade names, commercial products, or organizations imply endorsement by the U.S. Government.

## References

1. Vamathevan, J., et al., Applications of machine learning in drug discovery and development. Nature Reviews Drug Discovery, 2019. 18(6): p. 463–477.

2. Struble, T.J., et al., Current and Future Roles of Artificial Intelligence in Medicinal Chemistry Synthesis. Journal of Medicinal Chemistry, 2020. 63(16): p. 8667–8682.

3. Jiménez-Luna, J., et al., Artificial intelligence in drug discovery: recent advances and future perspectives. Expert Opinion on Drug Discovery, 2021: p. 1–11.

4. Strovel, J., et al., Early drug discovery and development guidelines: for academic researchers, collaborators, and start-up companies, in Assay Guidance Manual [Internet], S. Markossian, G. Sittampalam, and A. Grossman, Editors. 2012, Eli Lilly & Company and the National Center for Advancing Translational Sciences: Bethesda, MD.

5. Altekar, M., et al., Assay optimization: a statistical design of experiments approach. Clin Lab Med, 2007. 27(1): p. 139–54.

6. Fisher, R.A., Design of Experiments. British Medical Journal, 1936. 1(3923): p. 554.

7. Taylor, P.B., et al., Automated Assay Optimization with Integrated Statistics and Smart Robotics. Journal of Biomolecular Screening, 2000. 5(4): p. 213–225.

8. Shaw, R., et al., Overcoming Obstacles in the Implementation of Factorial Design for Assay Optimization. ASSAY and Drug Development Technologies, 2015. 13(2): p. 88–93.

9. Sittampalam, G.S., et al., Mapping biologically active chemical space to accelerate drug discovery. Nat Rev Drug Discov, 2019. 18(2): p. 83–84.

10. Godfrey, A.G., et al., A Perspective on Innovating the Chemistry Lab Bench. Front Robot AI, 2020. 7: p. 24.

11. Duncan, K.K., et al., Exploring Novel Biologically-Relevant Chemical Space Through Artificial Intelligence: The NCATS ASPIRE Program. Front Robot AI, 2019. 6: p. 143.

12. Michael, S., et al., A robotic platform for quantitative high-throughput screening. Assay Drug Dev Technol, 2008. 6(5): p. 637–57.

13. Ferreira, R.S., et al., Complementarity between a docking and a high-throughput screen in discovering new cruzain inhibitors. J Med Chem, 2010. 53(13): p. 4891–905.

14. Chen, J., et al., Selective and cell-active inhibitors of the USP1/ UAF1 deubiquitinase complex reverse cisplatin resistance in non-small cell lung cancer cells. Chem Biol, 2011. 18(11): p. 1390–400.

15. Coussens, N.P., et al., Assay Guidance Manual: Quantitative Biology and Pharmacology in Preclinical Drug Discovery. Clin Transl Sci, 2018. 11(5): p. 461–470.

16. Assay Guidance Manual [Internet], S.G. Markossian S, Grossman A, et al., Editor. 2004-, Eli Lilly & Company and the National Center for Advancing Translational Sciences: Bethesda, MD.

17. Zhang, J.H., T.D. Chung, and K.R. Oldenburg, A Simple Statistical Parameter for Use in Evaluation and Validation of High Throughput Screening Assays. J Biomol Screen, 1999. 4(2): p. 67–73.

18. Bitter, R., T. Mohiuddin, and M. Nawrocki, LabVIEW: Advanced programming techniques. 2006: CRC Press.

19. Inglese, J., et al., Quantitative high-throughput screening: a titration-based approach that efficiently identifies biological activities in large chemical libraries. Proc Natl Acad Sci U S A, 2006. 103(31): p. 11473–8.

20. Bar, H. and A. Zweifach, Z’ Does Not Need to Be > 0.5. SLAS DISCOVERY: Advancing the Science of Drug Discovery, 2020. 25(9): p. 1000–1008.

