## Supplemental Information for "Cross-platform Bayesian optimization system for autonomous biological assay development"

### Supplemental Table 1. Standardized historical cell-free papain assay protocol.

| Step | Parameter | Value | Description |
|---|---|---|---|
| 1a<br>1b | Reagent<br>Reagent | 3 µL<br>3 µL | Papain in assay buffer<br>Assay buffer |
| 2 | Compound | 17 nL | Compound addition |
| 3 | Time | 15 min | RT incubation |
| 4 | Reagent | 1 µL | Substrate (Z-FR-AMC) in assay buffer addition, reaction initiation |
| 5 | Time | 15 sec, 271 *g* | Centrifugation |
| 6 | Detection | $(E_x/E_m)$ = 340/460 nm | Fluorescence (Read 1) |
| 7 | Time | 10 min | Incubation (RT) |
| 8 | Detection | $(E_x/E_m)$ = 340/460 nm | Fluorescence (Read 2) |

| Step | Notes |
|---|---|
| 1a | Medium-binding black solid-bottom Greiner plates (789176-F). Dispense 3 µL of 1.33X papain in assay buffer (100 mM sodium acetate, pH 5.5, 5 mM DL-cysteine, 0.01% Tween-20 v/v) to columns 1, 2, 5-48 *via* Wako/Kalypsys dispenser. Papain was dispensed to produce 5 nM final concentration. Enzyme solution kept on ice during experiment. |
| 1b | Dispense 3 µL of assay buffer (100 mM sodium acetate, pH 5.5, 5 mM DL-cysteine, 0.01% Tween-20 v/v) to columns 3 and 4 *via* Wako/Kalypsys dispenser. Assay buffer solution kept on ice during experiment. |
| 2 | Compound addition by single pin-transfer (17 nL) using Wako pintool. Compound solutions in source plate: 169 nM to 10 mM DMSO stock solutions. Final concentrations: 0.664 nM to 39.2 µM, respectively. Final DMSO concentration: 0.5% (v/v). |
| 3 | Room temperature incubation. Kalypsys lids used as microplate coverings. Protected from light. |
| 4 | Dispense 1 µL of 4X substrate mixture in assay buffer (100 mM sodium acetate, pH 5.5, 5 mM DL-cysteine, 0.01% Tween-20 v/v) to columns 1-48 *via* Wako/Kalypsys dispenser. Z-FR-AMC substrate was dispensed to produce 2 µM final concentration. Substrate solution kept on ice during experiment. |
| 5 | Centrifugation step to remove bubbles. |
| 6 | ViewLux optics: Excitation filter 340 (30) nm, Emission filter 450 (20) nm. Energy = 2000. Exposure time = 2 s. |
| 8 | ViewLux optics: Excitation filter 340 (30) nm, Emission filter 450 (20) nm. Energy = 2000. Exposure time = 2 s. |

**Supplemental Table 2. Liquid handler protocol notes for autonomous and continuous papain biochemical assay optimization (Dispensers 1, 2) and plate washing (Dispenser 3).** Dispensers = Wako/Kalypsys/GNF, each with three heads.

| Dispenser | Head | # tips | Solution dispensed | Notes |
|---|---|---|---|---|
| 1 | 1 | 8 | Enzyme | Straight tips<br>3 μL protocol for each tip and volume scaled accordingly by expected weight<br>8 separate bottles, tubing lines, valve/tips, airlines, O-rings (each tip is set up with 1 of 8 concentrations of enzyme)<br>Dispense to columns 1, 2, 5-48<br>Motion profile: carriage return<br>Pre-dispense volume: 75 μL<br>1 cycle per microplate |
| | 2 | 1 | Buffer | Angled tip<br>3 μL protocol for each tip and volume scaled accordingly by expected weight<br>1 bottle, tubing line, valve/tip, airline, O-ring<br>Dispense to columns 3, 4<br>1 cycle per microplate |
| 2 | 2 | 8 | Substrate | Angled tip<br>1 μL protocol for each tip and volume scaled accordingly by expected weight<br>8 separate bottles, tubing, valve/tips, airlines, O-rings (each tip is set up with 1 of 8 concentrations of substrate)<br>Dispense to columns 1-48<br>Motion profile: serpentine<br>Pre-dispense volume: 50 μL<br>1 cycle per microplate |
| 3 | 1 | 8 | Wash (70% ethanol) | Straight tips<br>5 μL protocol using all 8 tips (for speed) and volume scaled accordingly by expected weight<br>2 separate bottles, 4 tubing lines, valve/tips, airlines, O-rings per bottle<br>Dispense to columns 1-48<br>Motion profile: serpentine<br>Aspiration height: set to when pins touch the bottom of the well<br>1 cycle per microplate |
| | 2 | 8 | Wash (RO/DI water) | Angled tips<br>6 μL protocol using all 8 tips (for speed) and volume scaled accordingly by expected weight<br>Lines hooked up directly to house RO/DI manifold (no refilling of bottles needed)<br>Dispense to columns 1-48<br>Motion profile: serpentine<br>Aspiration height: set to when pins touch the bottom of the well and speed is set accordingly to maximize evacuation of well contents (water at this point)<br>3 cycles per microplate |

**Supplemental Table 3. List of reference small-molecule inhibitors of papain enzymatic activity for validation studies.**

| Sample ID (NCATS) | Source | SMILES | LYCHI3 (NCATS) | logIC$_{50}$ [M], Offline | logIC$_{50}$ [M], Online |
|---|---|---|---|---|---|
| NCGC00389776-02, NCGC00093887-03 | BioVision, SigmaAldrich | CC(C)C[C@H](NC(=O)C1OC1C(O)=O)C(=O)NCCCCNC(N)=N | KBQAV24AD6N | -8.17 | -8.83 |
| NCGC00261865-01, NCGC00162359-04, NCGC00162359-02 | SigmaAldrich | CC1=CC=C(C=C1)[S+]([O-])(=O)N[C@@H](CCCCN)C(=O)CCl | HQ1JGUPA1TY | -5.52 | -5.94 |
| NCGC00260755-01, NCGC00015084-03 | SigmaAldrich | NC(=N)C1=CC=C(N)C=C1 | N6CT2MK5LSC | ND | -4.49 |
| NCGC00094419-06, NCGC00261842-01, NCGC00094419-05 | SigmaAldrich, NCGC SigmaAldrich | CC1=CC=C(C=C1)[S+]([O-])(=O)N[C@@H](CC2=CC=CC=C2)C(=O)CCl | KCBQLTXPGWS | ND | -4.49 |
| NCGC00261127-01, NCGC00015369-06 | SigmaAldrich | Cl\C1=C(/Cl)C2=C(C=CC=C2)C(=O)O1 | WFYQQ7V7NNN | > -4.30 | > -4.30 |
| NCGC00163460-01 | Biomol | CC(C)C[C@H](NC(C)=O)C(=O)N[C@@H](CC(C)C)C(=O)N[C@@H](CCCNC(N)=N)C=O | 7BR4SL7BG19 | -7.47 | -7.66 |
| NCGC00017341-01 | TimTec | CC(C)CC(NC(=O)C(NC(=O)NC(CC1=CC=CC=C1)C(O)=O)C2CCNC(=N)N2)C(=O)NC(CC3=CC=CC=C3)C=O | 12BM49DXS2V | -7.42 | -7.72 |
| NCGC00390338-01 | BioVision | CC(C)C(NC(=O)C(CCCN=C(N)N)NC(=O)NC(CC1=CC=CC=C1)C(O)=O)C(=O)NC(CCCN=C(N)N)C=O | AJZCDAXKJVD | -7.32 | -7.61 |
| NCGC00168270-01 | NCGC | FC1=CC=CC(NC2=NC(=NC(NC3CCCC3)=N2)C#N)=C1 | 2P5GL6C45RA | -6.82 | -6.75 |
| NCGC00049042-02 | Chemical Block | CCOC1=NC(=NC(=N1)N2CCCCC2)C#N | JL7M1KSPN9K | -6.52 | -6.65 |
| NCGC00168273-01 | NCGC | COC1=CC=CC(NC2=NC(=NC(NC3CCCC3)=N2)C#N)=C1 | 2RBHQDG7MQ4 | -6.52 | -6.88 |
| NCGC00168274-01 | NCGC | COC1=CC=C(NC2=NC(=NC(NC3CCCC3)=N2)C#N)C=C1 | DQ1Q2KY4N3A | -6.47 | -6.94 |
| NCGC00167649-01 | NCGC | N#CC1=NC(NC2CCCC2)=NC(NC3=CC=CC=C3)=N1 | FNYBXMGBXB5 | -6.47 | -6.90 |
| NCGC00168481-01 | NCGC | FC1=CC(NC2=NC(=NC(NC3CCCC3)=N2)C#N)=CC(F)=C1 | G9UTCJBFBAZ | -6.42 | -6.55 |
| NCGC00168271-01 | NCGC | FC1=CC=C(NC2=NC(=NC(NC3CCCC3)=N2)C#N)C=C1 | 9L6WQAQMQ4G | -6.42 | -6.85 |
| NCGC00168276-01 | NCGC | CC1=CC=CC(NC2=NC(=NC(NC3CCCC3)=N2)C#N)=C1 | 2HKWQP1ZV91 | -6.32 | -6.82 |
| NCGC00168272-01 | NCGC | ClC1=CC=CC(NC2=NC(=NC(NC3CCCC3)=N2)C#N)=C1 | 2RLBFNLC1RG | -5.92 | -6.31 |
| NCGC00015412-20 | Microsource | O=C1N([Se]C2=C1C=CC=C2)C3=CC=CC=C3 | HGKG9L7ZW8D | > -4.30 | > -4.30 |
| NCGC00389647-01 | APExBIO | NCC(=O)NCC(=O)NC(CC1=CC=C(O)C=C1)C(=O)NC(CCCN=C(N)N)C(O)=O | Q23NCTXQF22 | > -4.30 | -4.31 |
